# Role of human HSPE1 for OPA1 processing independent of HSPD1

**DOI:** 10.1101/2022.08.11.503680

**Authors:** Nelson Yeung, Daisuke Murata, Miho Iijima, Hiromi Sesaki

## Abstract

The mtHSP60/HSPD1-mtHSP10/HSPE1 system prevents protein misfolding and maintains proteostasis in the mitochondrial matrix. Altered activities of this chaperonin system have been implicated in human diseases, such as cancer and neurodegeneration. However, how defects in HSPD1 and HSPE1 affect mitochondrial structure and dynamics remains elusive. In the current study, we address this fundamental question in a human cell line, HEK293T. We found that the depletion of HSPD1 or HSPE1 results in fragmentation of mitochondria, suggesting a decrease in mitochondrial fusion. Supporting this notion, HSPE1 depletion led to proteolytic inactivation of OPA1, a dynamin-related GTPase that fuses the mitochondrial membrane. This OPA1 inactivation was mediated by a stress-activated metalloprotease, OMA1. In contrast, HSPD1 depletion did not induce OMA1 activation or OPA1 cleavage. These data suggest that HSPE1 controls mitochondrial morphology through a mechanism separate from its chaperonin activity.

## Introduction

Human mitochondria contain approximately 1,000-1,500 proteins, more than 99% encoded in nuclear DNA, synthesized in the cytosol, and imported to mitochondria (Pfanner et al., 2019). Only 13 proteins are encoded in mitochondrial DNA (Kopinski et al., 2021). Since the protein translocases that import proteins into mitochondria have small pores, nuclear-encoded mitochondrial proteins must be translocated into mitochondria in an unfolded conformation (Araiso et al., 2022). Once imported into mitochondria, proteins are folded, aided by chaperones via an oxidative protein folding pathway in the intermembrane space and an ATP-driven protein folding pathway in the matrix (Finka et al., 2016). After maturation, correct folding of mitochondrial proteins is ensured by chaperonins (Horwich and Fenton, 2020). Defects in mitochondrial chaperones and chaperonins have been linked to many human diseases (Bahr et al., 2022; Rodriguez et al., 2020).

In the mitochondrial matrix, two major types of machinery mediate protein folding: the mtHSP70-mtHSP40 system (chaperone) and mtHSP60-mtHSP10 system (chaperonin) (Bahr et al., 2022; Hayer-Hartl et al., 2016; Horwich and Fenton, 2020). The mtHSP70-mtHSP40 system, along with other proteins, facilitates client proteins to be correctly folded during import into the matrix through the translocase of the inner membrane (Bahr et al., 2022; Horwich et al., 1990; Shin et al., 2021; Song and Becker, 2022). The mtHSP60-mtHSP10 system counteracts protein misfolding and unfolding under physiological conditions and in response to different types of stress (Bahr et al., 2022; Song and Becker, 2022). The pioneering studies on the founding members of the mtHSP60-mtHSP10 system (the bacterial GroEL-GroES system) revealed that GroEL/mtHSP60 forms a barrel structure with two ring-shaped GroEL/mtHSP60 oligomers (Hayer-Hartl et al., 2016; Horwich and Fenton, 2020; Horwich et al., 1990). The GroEL/mtHSP60 barrel accommodates substrate proteins and mediates protein folding reactions using ATP hydrolysis. GroES/mtHSP10 binds to the top and bottom of the barrel and closes a lid to optimize reactions. GroES/mtHSP10 also regulates the cycle of ATP hydrolysis by GroEL/mtHSP60 and faciliate GroEL/mtHSP60 to bind and fold substrate proteins (Hayer-Hartl et al., 2016; Horwich and Fenton, 2020; Horwich et al., 1990). While GroEL and GroES are essential for bacterial cell viability, mtHSP60 and mtHSP10 are vital for mitochondrial function (Horwich et al., 1990).

Mitochondria are dynamic organelles and change their morphology through membrane fusion and division (Giacomello et al., 2020; Kraus et al., 2021; Murata et al., 2020a) (Friedman and Nunnari, 2014; Kashatus, 2018; Widlansky and Hill, 2018; Youle and van der Bliek, 2012). These dynamic processes play important roles in maintaining mitochondrial health (Itoh et al., 2013; Liesa and Shirihai, 2013; Roy et al., 2015; Serasinghe and Chipuk, 2017). For example, bioenergetic stress could lead to fragmentation of mitochondria due to decreased mitochondrial fusion relative to division, facilitating efficient mitophagy (Murata et al., 2020a). Three dynamin-related GTPases mediate mitochondrial fusion (Kameoka et al., 2018; Murata et al., 2020a). An inner membrane GTPase, OPA1, fuses the inner membrane through interactions with a mitochondria-specific phospholipid, cardiolipin (Kameoka et al., 2018). Inner membrane fusion is coupled to outer membrane fusion, which is controlled by two homologous GTPases, MFN1 and MFN2. A regulatory mechanism for mitochondrial fusion is proteolytic controls of these three GTPases (Giacomello et al., 2020; Kraus et al., 2021; Murata et al., 2020a). For example, OPA1 produces membrane-anchored long forms that mediate mitochondrial fusion, while its cleavage by a metalloprotease, OMA1, greatly inactivates OPA1’s ability to fuse the membrane (Ehses et al., 2009; Head et al., 2009; Kameoka et al., 2018; MacVicar and Langer, 2016; Murata et al., 2020b; Yamada et al., 2019; Yamada et al., 2018). MFN1 and MFN2 are subjected to degradation to dump their fusion activity via different ubiquitin E3 ligases and proteosomes (Dorn, 2020; Murata et al., 2020b; Yamada et al., 2019). Various stresses and bioenergetic deficits can activate these regulatory systems for mitochondrial fusion (Dorn, 2020; Giacomello et al., 2020; Kraus et al., 2021; Murata et al., 2020a). However, it remains unclear whether defects in the mtHSP60-mtHSP10 system induce changes in OPA1, MFN1, or MFN2.

In the current study, we depleted mtHSP60 and mtHSP10 in a human cell line and analyzed the impact on the three GTPases. We found that the knockdown of mtHSP60 or mtHSP10 induced fragmentation of mitochondria. To our surprise, mtHSP10 knockdown increased levels of inactive forms of OPA1 through activation of OMA1 without affecting MFN1 and MFN2 levels. In contrast, mtHSP60 knockdown did not affect OPA1, MFN1, or MFN2. These findings suggest that mtHSP10 and mtHSP60 affect mitochondrial morphology through multiple pathways independent of their chaperonin activities.

## Results

### Knockdown of HSPE1 enhances OPA1 processing independent of HSPD1

To understand the role of mitochondrial chaperones and chaperonins for mitochondrial fusion, we knocked down four matrix-located heat shock proteins — HSPD1 (mtHSP60), HSPE1 (mtHSP10), HSPA9 (mtHSP70), and DNAJA3 (mtHSP40) — using esiRNAs for 5 days in HEK293T cells. We then examined OPA1, MFN1, and MFN2 using Western blotting. Knockdown of each protein was confirmed by Western blotting of whole cell lysates except for HSPE1 (Fig. 1A and B). Since we could not detect HSPE1 by Western blotting using five commercially available HSPE1 antibodies (Fig. S1), we confirmed its knockdown by quantitative PCR (Fig. 1C). The lack of HSPE1 detection by Western blotting may be due to its small size (102 amino acids, 10,932 Da); when ectopically expressed in HEK293T cells, we were able to detect HSPE1-GFP (GFP consists of 238 amino acids, 27kDa) using two out of the five anti-HSPE1 antibodies on Western blotting, but not HSPE1-FLAG (FLAG consists of 8 amino acids, 1kDa) using anti-FLAG antibodies (Fig. S1). In addition, the expression level may also contribute to the efficiency of detection.

**Figure 1.**
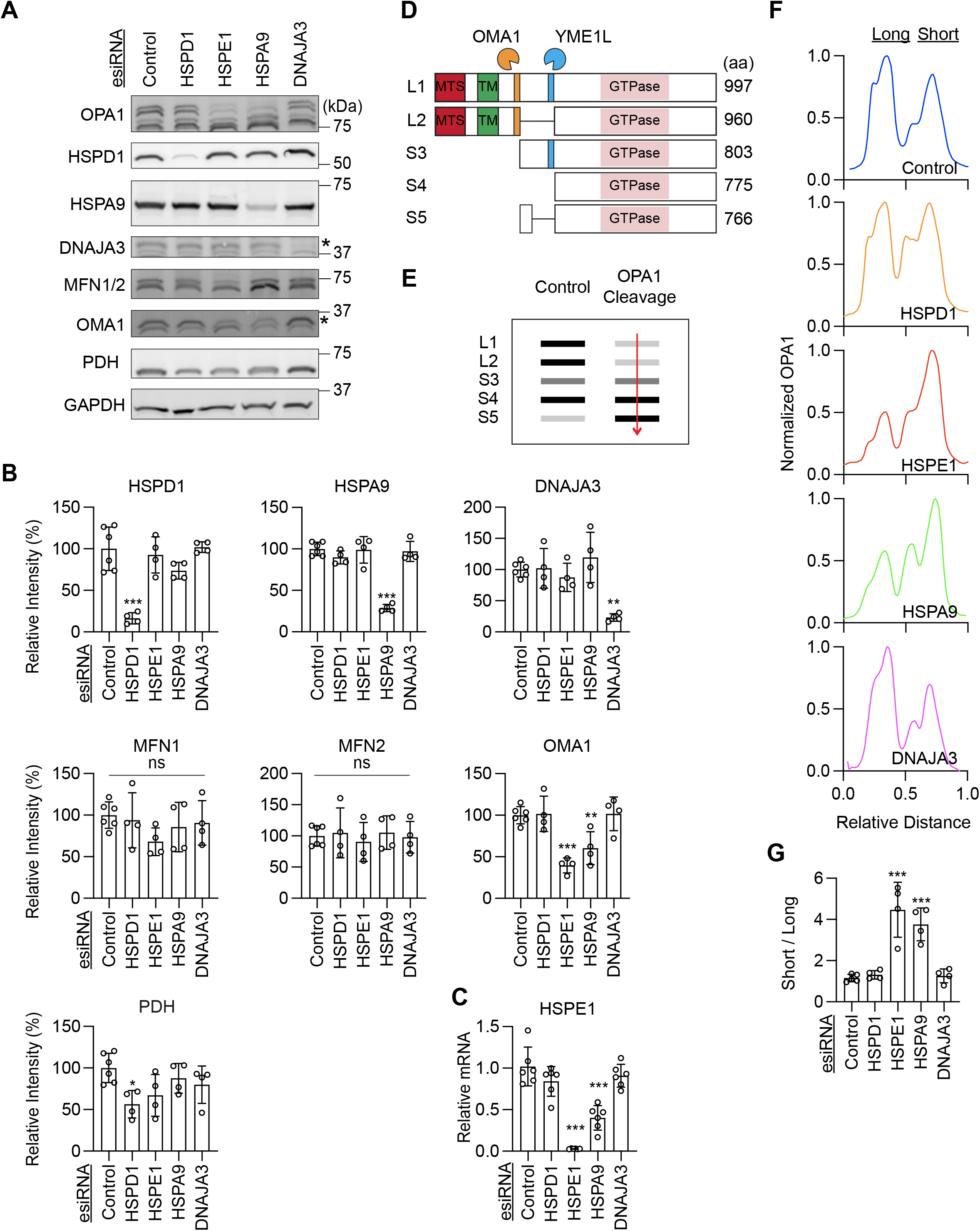
Knockdown of HSPE1 or HSPA9 induces OPA1 cleavage in HEK293T cells. A. The indicated mitochondrial chaperones and chaperonins were knocked down in HEK293T cells for 5 days using esiRNAs. Whole-cell lysates were analyzed by Western blotting with the indicated antibodies. The asterisks indicate the target band. B. Quantifications of band intensity normalized relative to GAPDH. Bars indicate averages ± SD (n=4 and 6). C. Relative mRNA levels of HSPE1 were determined by quantitative PCR in cells treated with the indicated esiRNAs. Bars indicate averages ± SD (n=6). D. Cleavage of OPA1 long isoforms by OMA1 and YME1L to produce short isoforms. E. OPA1 isoforms were quantified by measuring band intensity along a uniform line. F. A representative band intensity of OPA1 isoforms in cells with the indicated knockdown. G. The ratio of short isoforms of OPA1 over long isoforms is quantified (n=4). Statistical analysis was performed by one-way ANOVA with post hoc Tukey: *p <0.05, **p < 0.01, ***p < 0.001.

OPA1 is encoded by a single gene and generates multiple isoforms through alternative mRNA splicing and proteolytic processing (Kameoka et al., 2018; MacVicar and Langer, 2016; Murata et al., 2020b). In HEK293T cells, five isoforms, including two long forms (L1 and L2) and three short forms (S3, S4, and S5), were detected by Western blotting (Fig. 1A, D and E). S3 and S5 are produced via cleavage of L1 and L2, respectively, by the inner membrane metalloprotease OMA1 (Fig. 1D and E) (Ehses et al., 2009; Head et al., 2009; Kameoka et al., 2018; MacVicar and Langer, 2016; Murata et al., 2020b). However, in HEK293T cells (unlike in mouse embryonic fibroblasts, (Murata et al., 2020b), we noticed that the distances between L1 and L2 and between S4 and S5 in Western blots were sometimes too little to clearly separate them. Therefore, to analyze the conversion of L1 and L2 to S3 and S5, we measured the ratio of short forms relative to long forms (Fig. 1E). Interestingly, we found that the ratio of short forms over long forms decreased after the depletion of HSPE1 or HSPA9 (Fig. 1F and G). In contrast, we found no significant changes in the amounts of MFN1 or MFN2 after depletion of HSPD1, HSPE1, HSPA9, and DNAJA3 (Fig. 1A and B). Since the role of HSPA9 and DNAJA3 have been reported previously (Lee et al., 2015), we focused on HSPE1 in the current study.

HPSD1/mtHSP60 and HPSE1/mtHSP10 function together in protein folding in the mitochondrial matrix as chaperonins (Hayer-Hartl et al., 2016; Horwich and Fenton, 2020; Horwich et al., 1990). HSPD1 creates oligomeric rings and assembles into a barrel in which protein folding reactions occur using ATP binding and hydrolysis (Hayer-Hartl et al., 2016; Horwich and Fenton, 2020; Horwich et al., 1990). HSPE1 covers the apical sides of the HSPD1 barrel structure as a cap and regulates ATP hydrolysis. Both HSPD1 and HSPE1 are required for this protein folding mechanism in the matrix. Therefore, we were surprised by our finding that knockdown of HSPE1, but not HSPD1, stimulates the conversion of long OPA1 isoforms to short isoforms (Fig. 1F and G). These data suggest that HSPE1 has a role in OPA1 processing distinct from its protein folding function together with HSPD1. Alternatively, since the loss of HSPE1 could halt the function of HSPD1, the changes in OPA1 processing may be caused by a dominant negative effect of HSPD1 in the absence of HPSE1. This second model predicts that an additional knockdown of HSPD1 could mitigate the effect of HSPE1 knockdown on OPA1 processing. In the third model, if a decrease in protein folding function resulted in increased OPA1 processing, double knockdown of HSPD1 and HSPE1 may exacerbate enhanced OPA1 processing. To distinguish these models, we simultaneously knocked down HSPD1 and HSPE1 using esiRNAs (Fig. 2A and B). We found that additional knockdown of HSPD1 did not significantly change the OPA1 conversion in HSPE1-knockdown cells (Fig. 2C and D). These data suggest that HSPE1 depletions stimulate OPA1 processing independently of its chaperonin function and HSPD1.

**Figure 2.**
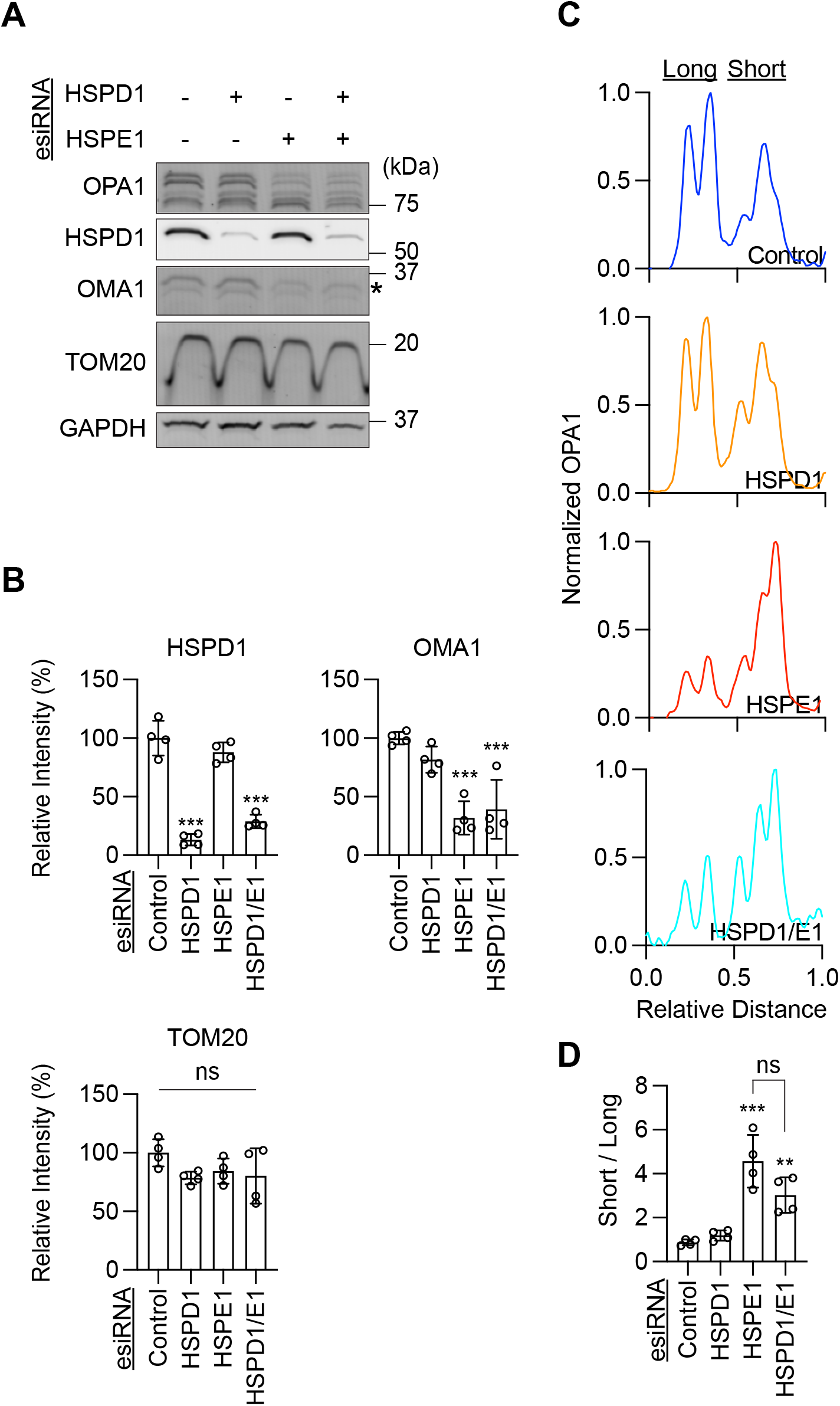
OPA1 cleavage induced by HSPE1 knockdown is independent of HSPD1. A. Western blot of HEK293T cells after individual or simultaneous knockdown of HSPD1 and HSPE1 using esiRNA. The asterisks indicate the target band. B. Quantitation of band intensity normalized relative to GAPDH. Bars indicate averages ± SD (n=4). C. A representative band intensity of OPA1 isoforms in cells with the indicated knockdown. D. The ratio of short isoforms of OPA1 over long isoforms is quantified (n=4). Statistical analysis was performed by one-way ANOVA with post hoc Tukey: **p < 0.01, ***p < 0.001.

### OMA1 mediates increased OPA1 processing in HSPEI-knockdown cells

OPA1 is cleaved by a metalloprotease located in the inner membrane, OMA1 (Acin-Perez et al., 2018; Baker et al., 2014; Ehses et al., 2009; Head et al., 2009; Kameoka et al., 2018; MacVicar and Langer, 2016; Murata et al., 2020b; Quiros et al., 2012; Rainbolt et al., 2016; Zhang et al., 2014). In response to mitochondrial stress, OMA1 is activated via its proteolytic cleavage followed by degradation. This stimulation mechanism assures a transient induction of its enzymatic activity (Ehses et al., 2009; Head et al., 2009; Kameoka et al., 2018; MacVicar and Langer, 2016; Murata et al., 2020b). After activation, OMA1 cleaves OPA1 long forms and generates its short forms. Therefore, decreased levels of OMA1 have been used as a read-out for its activation (Murata et al., 2020b). In HSPE1-knockdown cells, we found that OMA1 levels are decreased by Western blotting (Fig. 1A, 1B, 3A, and 3B). Based on these results, we hypothesized that OMA1 is activated upon HSPE1 depletion.

**Figure 3.**
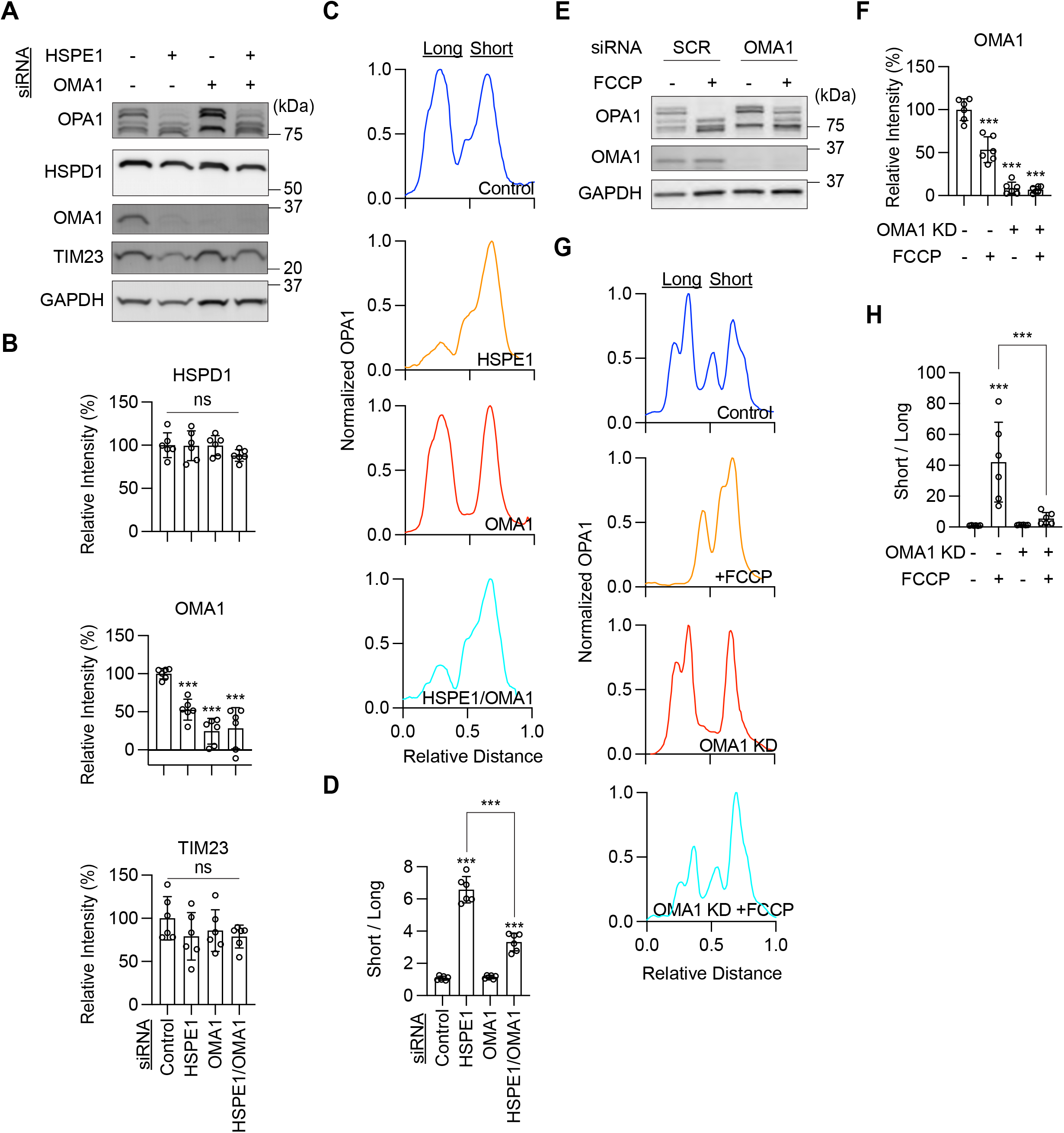
Knockdown of OMA1 shows a reduction in OPA1 cleavage induced by HSPE1 knockdown. A. Western blotting of HEK293T cells depleted for HSPE1 and OMA1 using siRNAs. B. Quantitation of band intensity. Bars indicate averages ± SD (n=6). C. A representative band intensity of OPA1 isoforms in cells with the indicated knockdown. D. Quantifying OPA1 isoforms using the ratio of short isoforms over long isoforms in C (n=6). E. Western blot with or without OMA1 siRNA knockdown in the presence or absence of 10 μM FCCP treatment with the indicated antibodies. F. A representative band intensity of OPA1 isoforms in cells with the indicated knockdown. D. The ratio of short isoforms of OPA1 over long isoforms is quantified (n=4). Statistical analysis was performed by one-way ANOVA with post hoc Tukey: **p < 0.01, ***p < 0.001.

To test whether OMA1 is responsible for converting OPA1 long forms to short forms in HSPE1 depleted cells, we knocked down OMA1 along with HSPE1 using siRNAs in HEK293T cells (Fig. 3A and B). Western blotting of whole cell lysates showed that OPA1 knockdown partially rescued the ratio of the short forms over the long forms (Fig. 3C and D). As a control experiment, we treated HEK293T cells with a proton ionophore, carbonyl cyanide 4-(trifluoromethoxy)phenylhydrazone (FCCP), which dissipates the membrane potential across the inner membrane. FCCP treatments activate OMA1 and induce OPA1 processing (Murata et al., 2020b). We found that FCCP treatments decrease OMA1 levels and that OMA1 knockdown partially rescues FCCP-induced OPA1 processing (Fig. 3E, F and G). These data, taken together, suggest that the loss of HSPE1 activates OMA1 and induces proteolytic cleavage of OPA1 from long forms to short forms.

### Mutations in the mobile loop inhibit the role of HSPE1 in OPA1 processing

The structure of the bacterial homolog of HSPD1 (GroEL) together with HSPE1 (GroES) has been solved (Hayer-Hartl et al., 2016; Horwich and Fenton, 2020; Hunt et al., 1996; Xu et al., 1997). Oligomeric GroEL rings form barrel structures, and GroES caps the top and bottom of barrels. A mobile loop in GroES with 20 amino acids mediates its interaction with the apical region of the GroEL barrels. Mutations in three amino acids (Ile 25, Val 26, and Leu 27) in bacterial GroES have been shown to block its binding to HSPD1 and protein folding (Nojima et al., 2012; Xu et al., 1997). The amino acid sequence of the mobile loop is conserved in bacterial GroES and human HSPE1 (Fig. 4A). (Bie et al., 2016). To test the functional importance of the mobile loop, we mutated the corresponding three residues (Ile 31, Met 32, Leu 33) to alanine in a siRNA-resistant version of HSPE1. WT and AAA mutant HSPE1 proteins fused to GFP were expressed in HSPE1-knockdown cells. We found that WT HSPE1-GFP rescued increased OPA1 cleavage in HSPE1-knockdown cells (Fig. 4B–D). In contrast, the AAA mutant HSPE1-GFP failed to do so. These data demonstrate that the mobile loop is important for the function of HSPE1 in the control of OPA1 processing. Since HSPD1 is not involved in the enhanced OPA1 processing observed in HSPE1-knockdown cells (Fig. 4B–D), the mobile loop has a critical role that is separable from its interaction with HSPD1 and protein folding.

**Figure 4.**
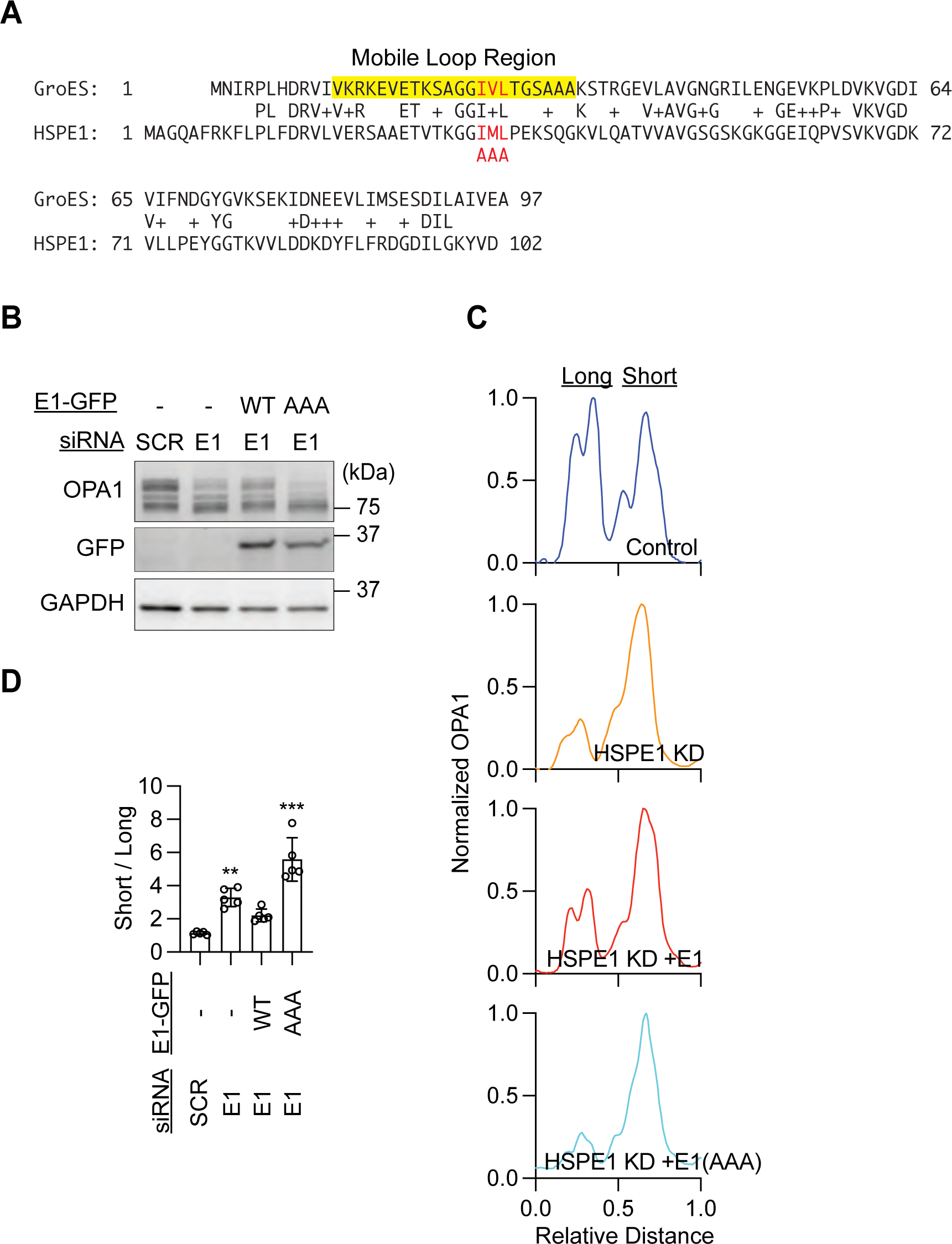
HSPE1 carrying mutations in the mobile loop fails to rescue increased OPA1 processing in HSPE1-knockdown cells. A. Amino acid sequence of bacterial GroES and human HSPE1. The mobile loop is highlighted in yellow. Mutated residues are indicated in red. B. HSPE1-knockdown HEK293T cells were transfected with siRNA-resistant forms of HSPE1 fused GFP. WT and mutant HSPE1-GFP carrying the AAA mutations are used. C. A representative band intensity of OPA1 isoforms in cells with the indicated knockdown. D. Quantifying OPA1 isoforms using the ratio of short isoforms over long isoforms (n=5). Significance was calculated using one-way ANOVA with post hoc Tukey: ** p < 0.01, ***p < 0.001.

### Fragmentation of mitochondria is induced by the depletion of mitochondrial chaperones and chaperonins

To test the role of HSPD1 and HSPE1 for mitochondrial morphology, we examined mitochondria using laser scanning confocal immunofluorescence microscopy using antibodies to TOM20 (a mitochondrial outer membrane protein) and pyruvate dehydrogenase (PDH, a matrix protein) in HEK293T cells knocked down for HSPD1 and HSPE1 as well as HSPA9 and DNAJA3. We found that individual knockdown of all four proteins produced shorter mitochondria (Fig. 5A and B). These data show that these proteins are important for maintaining mitochondrial morphology. Since the ratio of OPA1 short forms and long forms was only altered in HSPE1- or HSPA9-knockdown cells, mitochondrial fragmentation in HSPD1- or DNAJA3-knockdown cells likely resulted from other mechanisms. To test whether a decrease in mitochondria size in HSPE1-knockdown cells depends on OMA1, we simultaneously knocked down HSPE1 and OMA1 in HEK293T cells. We found that the OMA1 depletion modestly increased the size of mitochondria in HSPE1-knockdown cells (Fig. 6A and B). These data suggest that mitochondrial fragmentation is induced not only by OMA1-mediated OPA1 processing but also by additional mechanisms.

**Figure 5.**
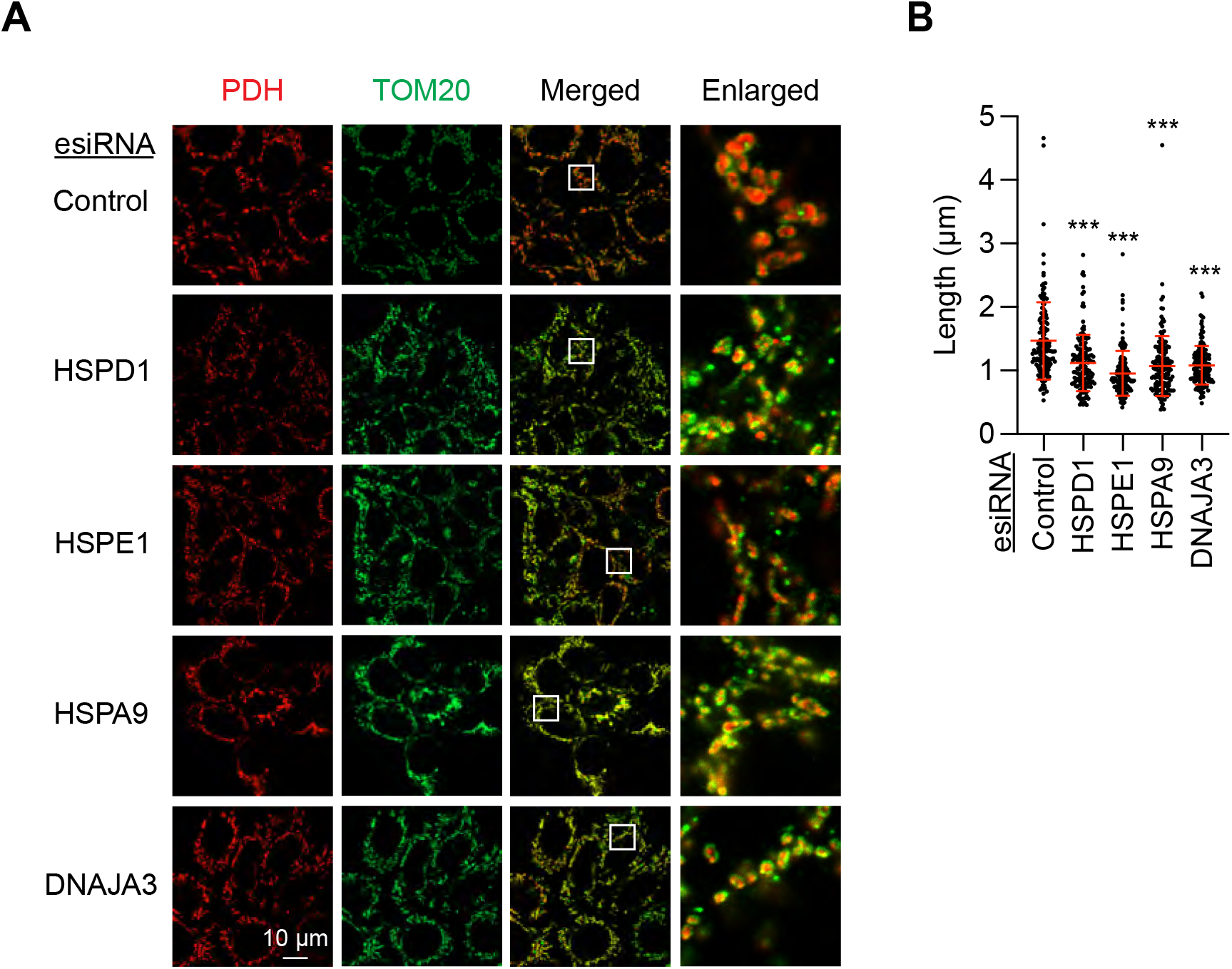
Mitochondria length decreases with the knockdown of mitochondrial chaperones and chaperonins. A. Confocal immunofluorescence microscopy of the indicated knockdown HEK293T cells using antibodies to pyruvate dehydrogenase (PDH, a matrix protein) and TOM20 (an outer membrane protein). Boxed regions are enlarged. B. Quantitation of mitochondria length using TOM20 staining. Bars indicate averages ± SD (n=150, 150 mitochondria in 30 cells were quantified for each knockdown). Significance was calculated using one-way ANOVA with post hoc Tukey: ***p < 0.0001.

**Figure 6.**
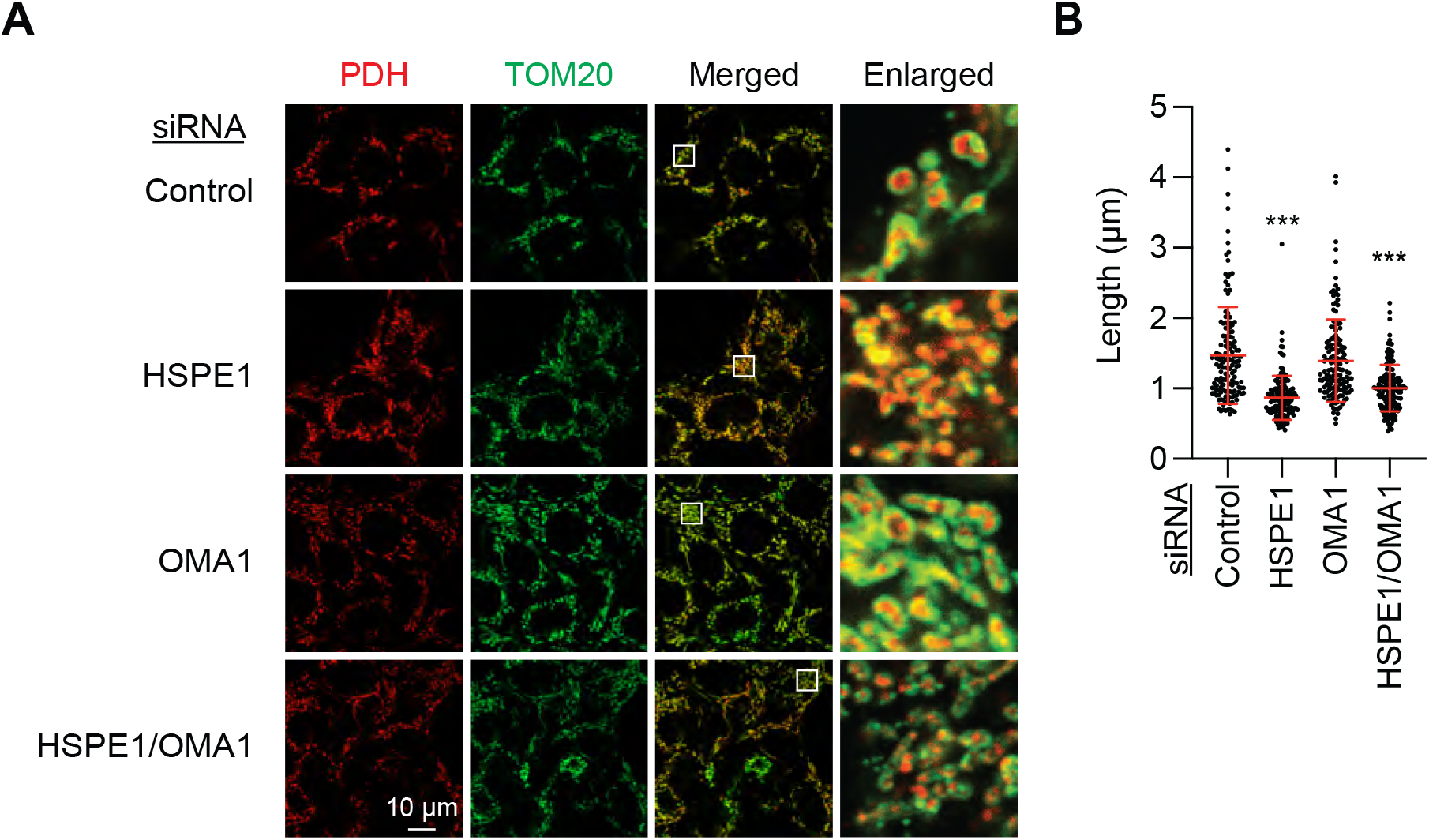
Mitochondria length was not rescued by the depletion of OMA1 in HSPE1-knockdown cells. A. Confocal immunofluorescence microscopy of the indicated knockdown HEK293T cells using antibodies to PDH and TOM20. Boxed regions are enlarged. B. Quantitation of mitochondria length using TOM20 staining. Bars indicate averages ± SD (n=150, 150 mitochondria in 30 cells were quantified for each knockdown). Significance was calculated using one-way ANOVA with post hoc Tukey: ***p < 0.001.

## Discussion

Here, we report that the loss of HSPE1 (mtHSP10) leads to the activation of the stress-responsive metalloprotease OMA1 and promotes the cleavage of OPA1. In contrast, the depletion of HSPD1 (mtHSP60), a functional partner of HSPE1 in protein folding, did not induce OMA1 activation or OPA1 cleavage. These data suggest that HSPE1 plays a role in protection from mitochondrial stress independently of HSPD1. At this moment, we do not know how the loss of HSPE1 activates OMA1. If HSPE1 depletion directly activates OMA1, one possibility is that a pool of HSPE1 is associated with OMA1 as a negative regulator, suppressing OMA1 activation at the steady state. In this model, dissociation of HSPE1 from OMA1 can activate this metalloprotease. However, our immunoprecipitation experiments found no stable association between HSPE1 and OMA1.

OMA1 is activated by various stressors, such as reactive oxygen species (Ehses et al., 2009; Head et al., 2009; Kameoka et al., 2018; MacVicar and Langer, 2016; Murata et al., 2020b). Given that HSPE1 deficiency induces OMA1 activation, some mitochondrial stress may promote OMA1 activation through changes in HSPE1 levels or function. Indeed, a previous study has suggested that sequestration of HSPE1 by pathogenic alpha-synuclein in the cytosol results in mitochondrial dysfunction and contributes to Parkinson’s disease (Szego et al., 2019). Parkinson’s disease has been strongly associated with aberrant mitochondrial structure and function (Panicker et al., 2021). In future studies, it would be interesting to test if alpha-synuclein accelerates OPA1 conversion.

In addition to the chaperonin function of HSPD1 in mitochondria, previous studies have also reported the roles of HSPD1 outside mitochondria in protein degradation and TNF-alpha-mediated activation of the IKK/NF-kappaB survival pathway in the cytosol (Choi et al., 2015; Chun et al., 2010; Kalderon et al., 2015). Interestingly, HSPD1 is further exported from cells and regulates the innate and adaptive immune system (Grundtman et al., 2011). Like HSPD1, HSPE1 might also play non-chaperonin roles inside or outside mitochondria or even in the extracellular space beyond protein folding.

Mitochondria are equipped with various mechanisms for stress response and quality control to protect their bioenergetic, metabolic, and signaling functions (Eldeeb et al., 2022; Murata et al., 2020a; Sugiura et al., 2014). For example, superoxide dismutases and glutathione peroxidases convert toxic reactive oxygen species to non-toxic molecules (Murphy et al., 2022). Intramitochondrial proteases degrade misfolded or unfolded protein. In addition, mitochondrial-derived vesicles and mitophagy sense damage to mitochondria, isolate damaged regions, remove such damaged portions from the rest of the mitochondria or even eliminate the whole mitochondria (Eldeeb et al., 2022; Murata et al., 2020a; Sugiura et al., 2014). Our finding of a role for HSPE1 independent of protein folding for OMA1 activation and OPA1 processing may add another layer of defense mechanisms that maintain mitochondrial health against physiological changes and pathological insults. It would be of great interest to define the underlying molecular mechanism by which HSPE1 senses and eliminates damage in mitochondria.

## Materials and Methods

### Cells and plasmids

HEK293T cells were cultured in Dulbecco’s Modified Eagle Medium containing 10% fetal bovine serum. esiRNAs and Silencer Select siRNAs were obtained from Millipore Sigma and Thermo Fisher, respectively (esiRNA: HSPD1 [EHU113521], HSPE1 [EHU112631], HSPA9 [EHU011841], DNAJA3 [EHU091601]; siRNA: Negative control [4390843], HSPD1 [S7003], HSPE1 [S7005], OMA1 [S41775]). These RNAs were transfected into HEK293T using Lipofectamine RNAiMAX (13778150, Themo Fisher). Human HSPE1-GFP was cloned into pcDNA3.1 from the HSPE1 (NM_002157.2) ORF Clone (OHu17870D, Genscript). The HSPE1-GFP constructs were transfected into cells using Lipofectamine 2000 (11668019, Themo Fisher).

### Western blotting

Cells were harvested and lysed in RIPA buffer (9806S, Cell Signaling Technology) supplemented with cOmplete Mini EDTA-free Protease Inhibitor Cocktail (11836170001, Roche) on ice (Murata et al., 2020b). The lysates were centrifuged at 16,000 g for 10 min at 4°C, and the supernatants were collected. Proteins were separated by SDS-PAGE and transferred onto Immobilon-FL Transfer Membrane (Millipore). The membranes were blocked in PBS-T (PBS containing 0.05% Tween 20) containing 3% BSA at room temperature for 1 h and then incubated with primary antibodies in PBS-T containing 3% BSA at 4°C overnight. The antibodies used were HSPD1 (1:1,000 dilution, 12165, Cell Signaling), DNAJA3 (1:1,000 dilution, 11088-1-AP; Proteintech), HSPA9 (1:5,000 dilution, 14887-1-AP; Proteintech), OPA1 (1:1,000 dilution, 612607; BD Biosciences), GAPDH (1:1,000 dilution, MA5-15738; Invitrogen), TOM20 (1:2,000 dilution, sc-11415; Santa Cruz Biotechnology), OMA1 (1:5,000 dilution, sc-515788; Santa Cruz Biotechnology), MFN1/2 (1:1,000 dilution, ab57602; Abcam), PDH (1:1,000 dilution, ab110338; Abcam), TIM23 (1:5,000 dilution, 611223; BD), and GFP (1:5,000 dilution, a gift from Dr. Peter Devreotes) (Adachi et al., 2020). The membranes were washed three times in PBS-T, followed by incubation with fluorescently labeled appropriate secondary antibodies at room temperature for 1 h. After washing the membranes three times in PBS-T, fluorescence signals were detected using a Typhoon laser-scanner platform (Amersham). Band intensity was quantified using the NIH FIJI software (Murata et al., 2020b).

### Real-time quantitative PCR

Total RNAs were purified from HEK293T cells using an RNeasy Mini Kit (74106; Qiagen) and reverse-transcribed using a ReadyScript cDNA Synthesis Mix (RDRT; Sigma-Aldrich) (Yamada et al., 2018; Yamada et al., 2022). PCR was performed using Quant Studio 3 (Thermo Scientific) and PowerUp SYBR Green Master Mix (A25741; Thermo Scientific). The DNA oligos are CAGTAGTCGCTGTTGGATCG and TCCAACTTTCACGCTAACTGG.

### Immunofluorescence microscopy

HEK293T cells were fixed in pre-warmed (37°C) PBS containing 4% paraformaldehyde for 20 min, washed three times in PBS, permeabilized with PBS containing 0.1% Triton X-100 for 8 min, washed again three times in PBS, and blocked in PBS containing 0.5% BSA at room temperature for 30 min (Adachi et al., 2016; Murata et al., 2020b). Cells were then incubated with antibodies to PDH (1:400 dilution) and TOM20 (1:300 dilution) at 4°C overnight. Cells were washed three times in PBS and incubated with Alexa 488-conjugated anti-mouse IgG (A21202, Thermo Fisher Scientific) and Alexa 568-conjugated anti-rabbit IgG (A10042, Thermo Fisher Scientific) at room temperature for 1 h. Cells were washed again, three times in PBS. The samples were observed using an LSM800 GaAsP laser scanning confocal microscope with a 100× objective (Murata et al., 2020b). To measure the mitochondria length, a circle with a 10 μm diameter was drawn on each cell, and the length of at least five mitochondria was measured in the long axis based on an anti-TOM20 signal using the NIH FIJI software (Adachi et al., 2020).

## Acknowledgments

We are grateful to past and present members of the Iijima and Sesaki labs for helpful discussions and technical assistance. This work was supported by NIH grants to MI (GM131768) and HS (GM144103).

## Author Contributions

NY, MI, and HS conceived the project and designed the study. NY, DM, and MI performed experiments and analyzed data. NY, MI, and HS wrote the manuscript.

---

**Figure S1.**
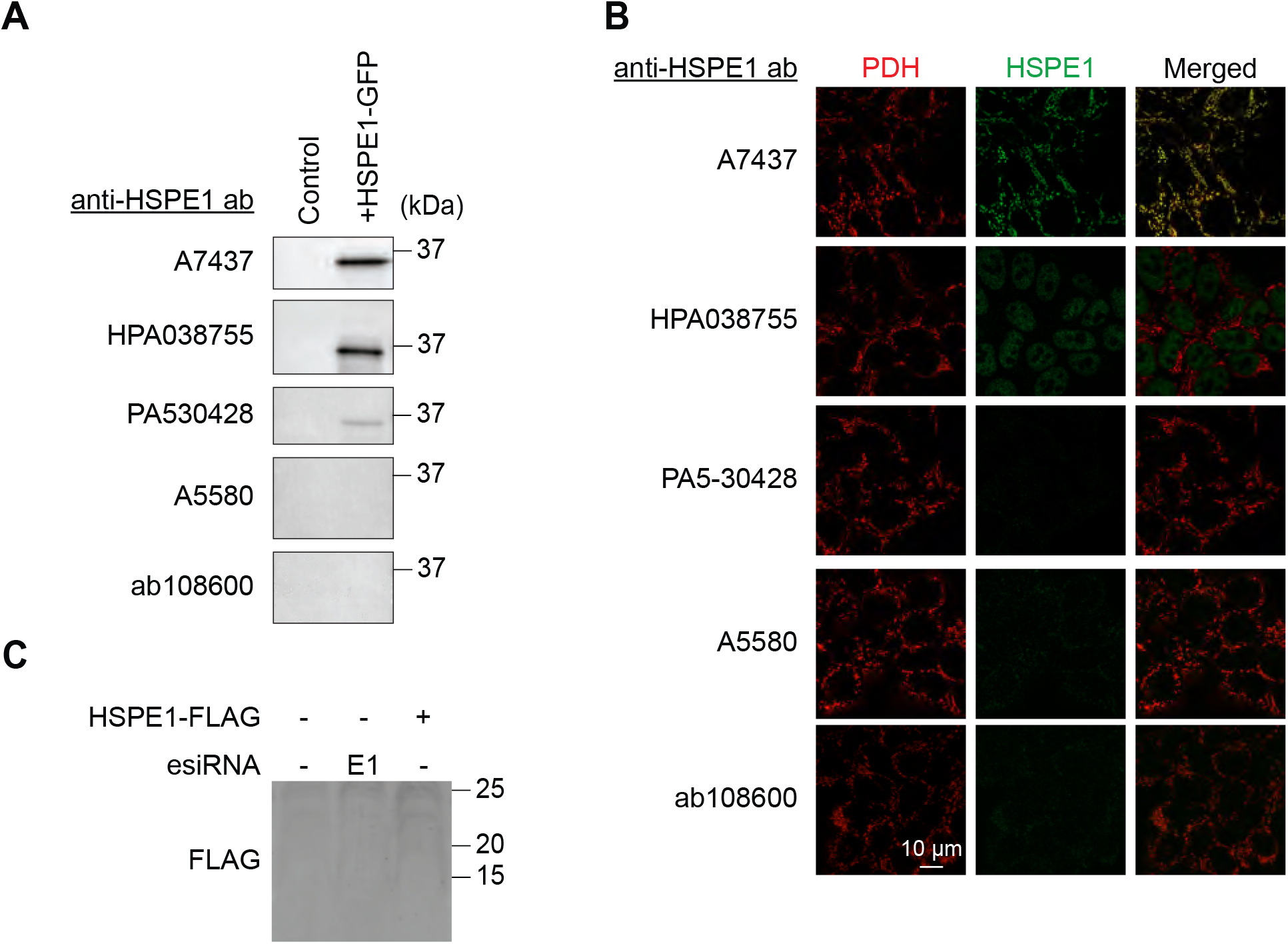
Analysis of anti-HSPE1 antibodies. A. HEK293T cells were transfected with HSPE1-GFP and analyzed by Western blotting with anti-HSPE1 antibodies (Abclonal [A7437], Sigma-Aldrich [HPA038755], Invitrogen [PA5-30428], Abclonal [A5580], and Abcam [ab108600]). B. HEK293T cells were subjected to immunofluorescence microscopy using antibodies to PDH and HSPE1. C. Western blotting of HEK293T cells depleted for HSPE1 or transfected with HSPE1-FLAG using anti-FLAG antibodies (Sigma [F7425]).

